# Nationwide population genetic screening improves outcomes of newborn screening for hearing loss in China

**DOI:** 10.1101/502088

**Authors:** Qiuju Wang, Jiale Xiang, Jun Sun, Yun Yang, Jing Guan, Dayong Wang, Cui Song, Ling Guo, Hongyang Wang, Yaqiu Chen, Junhong Leng, Xiaman Wang, Junqing Zhang, Bing Han, Jing Zou, Chengbin Yan, Lidong Zhao, Hongyu Luo, Yuan Han, Wen Yuan, Hongyun Zhang, Wei Wang, Jian Wang, Huanming Yang, Xun Xu, Ye Yin, Cynthia C. Morton, Lijian Zhao, Shida Zhu, Jun Shen, Zhiyu Peng

**Affiliations:** Department of Otolaryngology-Head and Neck Surgery, Chinese PLA Institute of Otolaryngology, Chinese PLA General Hospital, Beijing 100853, China; BGI Genomics, BGI-Shenzhen, Shenzhen 518083, China; Tianjin Medical Laboratory, BGI-Tianjin, BGI-Shenzhen, Tianjin 300308, China; Binhai Genomics Institute, BGI-Tianjin, BGI-Shenzhen, Tianjin 300308, China; Brigham and Women’s Hospital, Harvard Medical School, Boston, MA 02115, USA; Children’s Hospital of Chongqing Medical University, Chongqing 400014, China; Jining Maternal and Child Health Care Hospital, Jining 272000, China; Tianjin Women and Children’s Health Centre, Tianjing 300308, China; BGI Clinical Laboratory, BGI-Shenzhen, Shenzhen 518083, China; MGI, BGI-Shenzhen, Shenzhen 518083, China; Wuhan BGI Clinical Laboratory, BGI-Shenzhen, Wuhan 430074, China; BGI-Beijing, BGI-Shenzhen, Beijing 101300, China; BGI-Shenzhen, Shenzhen 518083, China; James D. Watson Institute of Genome Sciences, Hangzhou 310058, China; China National GeneBank, BGI-Shenzhen, Shenzhen 518120, China; University of Manchester, Manchester M13 9PL, UK; Shenzhen Engineering Laboratory for Innovative Molecular Diagnostics, Shenzhen 518120, China

**Keywords:** genetic screening, hearing screening, hearing loss, newborns, clinical benefits

## Abstract

**Purpose:** Concurrent newborn hearing and genetic screening has been reported, but its benefits have not been statistically proven due to limited sample sizes and outcome data. To fill this gap, we analyzed outcomes of a large number of newborns with genetic screening results.

**Methods:** Newborns in China were screened for 20 hearing-loss-related genetic variants from 2012–2017. Genetic results were categorized as positive, at-risk, inconclusive, or negative. Hearing screening results, risk factors, and up-to-date hearing status were followed-up via phone interviews.

**Results:** We completed genetic screening on one million newborns and followed up 12,778. We found that a positive genetic result significantly indicated a higher positive predictive value of the initial hearing screening (60% vs. 5.0%, P<0.001) and a lower rate of loss-to-follow-up (5% vs. 22%, P<0.001) than an inconclusive one. Importantly, 42% of subjects in the positive group with reported or presymptomatic hearing loss were “missed” by conventional hearing screening. Furthermore, genetic screening identified 0.23% of subjects predisposed to preventable ototoxicity.

**Conclusion:** Our results demonstrate that limited genetic screening identified additional cases, reduced loss-to-follow-up, and informed families of ototoxicity risks, providing convincing evidence to support integrating genetic screening into universal newborn hearing screening programs.

## Introduction

Hearing loss is one of the most common birth defects with life-long impact that may be ameliorated by early detection and intervention.^1^ Newborn hearing screening (NHS) programs have been globally adopted since the 1990s. They serve as a primary approach to detecting hearing loss in neonates and triggering early intervention.^2^ As designed, conventional hearing screening does not detect mild hearing loss nor offer prognosis to delayed-onset or drug-induced hearing loss.^2–5^ Hearing screening is unable to elucidate the etiology of hereditary hearing loss, which accounts for at least 50% to 60% of childhood hearing loss.^5^ Furthermore, only 10–15% of refers from hearing screening are confirmed to have permanent hearing loss.^5^ Low positive predictive values (PPV) may contribute to high rates of loss to follow-up (LFU) after the initial hearing screening,^6^ which defeats the purpose of NHS.

Limited genetic screening of a small number of genes commonly associated with hearing loss (*GJB2*, *SLC26A4* and *MT-RNR1*) to improve the detection of late-onset prelingual hearing loss was firstly proposed in 2006.^5^ Targeted screening of newborns for pathogenic variants in these genes represents an affordable paradigm. Studies have demonstrated the feasibility of concurrent hearing and genetic screening.^7–11^ However, the benefits of such practices have not been quantified due to limited sample sizes and outcome data. Herein we report results of genetic screening on ∼1.2 million newborns in China and outcomes of 12,778 infants with genetic findings followed-up via phone interviews.

## Methods

### Study design

This study was performed with the approval of the Institutional Review Board of BGI.

We enrolled newborns nationwide in China who received genetic screening from March 2012 to September 2017. Families or guardians of subjects with at least one variant identified between March 2012 and December 2016 were targeted for follow-up phone interviews from November 2016 to March 2017 (first interview period). Given the relatively small number of subjects with biallelic variants, we interviewed additional subjects with biallelic autosomal or homoplasmic mitochondrial variants identified from January to September 2017 between December 2017 and January 2018 (second interview period) to increase the power of the study (Figure 1). Subjects with biallelic variants but reported no hearing loss during the first interview period were recontacted during the second interview period. Newborns without any variant were not followed up because clinical management remains the same as the standard of care (**Supplementary Figure S1** online).

**Figure 1.**
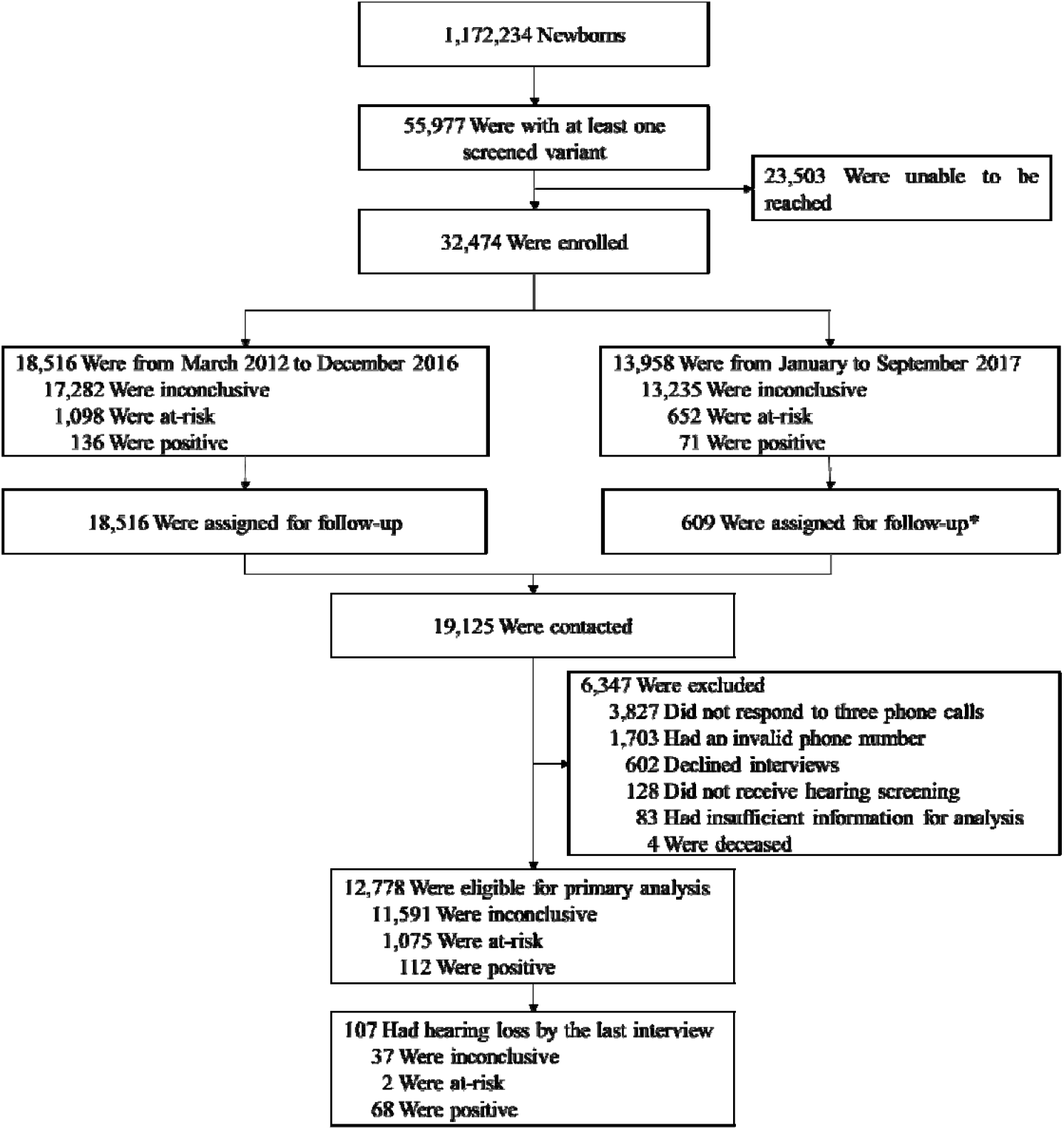
Enrollment and outcomes of subjects par in follow-up interviews. NHS denotes newborn hearing screening. Genetic results were classified as follows: positive (biallelic either *GJB2* or *SLC26A4*), inconclusive (a heterozygo in *GJB2* or *SLC26A4*, or presence of any *GJB3* varia (any mitochondrial variant), and negative (no variant). * Considering limited samples size of subjects wit variants from 2012–2016, we initiated interviews group of subjects with biallelic autosomal or hom mitochondrial variants from January to Septem including subjects with positive (n = 71), homoplasm (n = 536) and homozygous *GJB3* (n = 2) results were ta follow-up.

### Newborn genetic screening

The limited genetic screening entailed genotyping 20 variants in *GJB2*, *SLC26A4*, *MT-RNR1* and *GJB3* (**Supplementary Table S1** online) by MALDI-TOF mass spectrometry as previously validated and reported.^7,9^ Genetic screening results were classified as follows: positive (biallelic variants in either *GJB2* or *SLC26A4*), inconclusive (a heterozygous variant in *GJB2* or *SLC26A4*, or presence of any *GJB3* variant), at-risk (any mitochondrial variant), and negative (none of the 20 screened variants identified). All homozygous results were Sanger confirmed.

Each subject was catogoried into one and only one group based on the genetic results. When a subject’s results fell into more than one category, the subject was assigned to a group in the following order of priority: positive, at-risk, inconclusive, and negative.

The cost of genetic screening differed by regions, ranging from 32 to 48 US dollars. The turnaround time was around five working days upon receipt of a newborn’s dried blood spot specimen. Results were generally available by two to four weeks of age.

### Newborn hearing screening

Newborn hearing screening results were collected via phone interviews. The initial hearing screening in China was typically conducted by a trained nurse in the birth hospital 48–72 hours after birth using otoacoustic emission (OAE). Newborns who do not pass the initial NHS are referred for re-test around 42 days of age when OAE and/or automated auditory brainstem response (AABR) tests are performed. Newborns who do not pass the 42-day re-test are referred for comprehensive diagnostic audiometry conducted by audiologists generally around three months of age (**Supplementary Figure S1** online).

### Phone interview

We followed up newborns with a non-negative genetic result via phone interviews, during which we collected self-reported hearing screening results, hearing loss risk factors, and up-to-date hearing status. Subjects were considered LFU, if they were referred from the initial inpatient NHS but did not take either the 42-day re-test or the three-month audiologic evaluation. A subject was considered to have hearing loss: 1) if a formal audiologic evaluation determined a hearing threshold above 25 decibels, 2) if hearing rehabilitation had been implemented (*e.g*., hearing aids and cochlear implants), or 3) if hearing loss was reported based on behavioral assessment. The severity of hearing loss was graded according to the World Health Organization’s standards for children, namely profound (over 81 decibels), severe (61∼80 decibels), moderate (31∼60 decibels), and mild/slight (26∼30 decibels).^12^ The degree of hearing loss was “unspecified” if audiologic evaluation results were not available.

### Statistical analysis

Subjects with sufficient information obtained from phone interviews were included for the primary analysis. For categorical data, summary data were reported as frequencies and percentages, and chi-square tests were used for between-group comparisons. Age was reported as means ± standard deviations (SD), medians, and ranges in months. The difference in age between *GJB2* and *SLC26A4* positive subgroups was tested using the Mann-Whitney U test. Confidence intervals were computed using the Clopper-Pearson method. A P value of less than 0.05 was considered statistically significant. Statistical analysis was performed with IBM SPSS Statistics, version 24 (SPSS).

## Results

### Study population

From 2012 to 2017, we successfully conducted genetic screening of 1,172,234 newborns, finding 360 (0.03%) positive, 2,638 (0.23%) at-risk, 52,979 (4.52%) inconclusive, and 1,116,257 (95.22%) negative results (**Supplementary Table S2** online).

To evaluate outcomes of genetic screening, we identified 55,977 subjects (4.78%) harboring at least one of the screened variants (Figure 1), including 32,474 subjects with contact information for phone interviews. We contacted 19,125 subjects from both interview periods combined, and 12,778 (66.81%; mean age = 8.1±5.7 months, median age = 6 months, range 2–56 months) were eligible for the primary analysis, including 112 (0.9%) positive, 11,591 (90.7%) inconclusive, and 1,075 (8.4%) at-risk for ototoxicity. By the time of the last interviews, 107 (0.8%) subjects were reported to have hearing loss (Figure 1).

Table 1 shows the demographic characteristics of the 12,778 subjects. About 79.0% were younger than one year old, and 54.2% were male. Most subjects (86.9%) reported no risk factor for hearing loss, and 1,177 (9.2%) infants had been admitted to a neonatal intensive care unit.

**Table 1.**
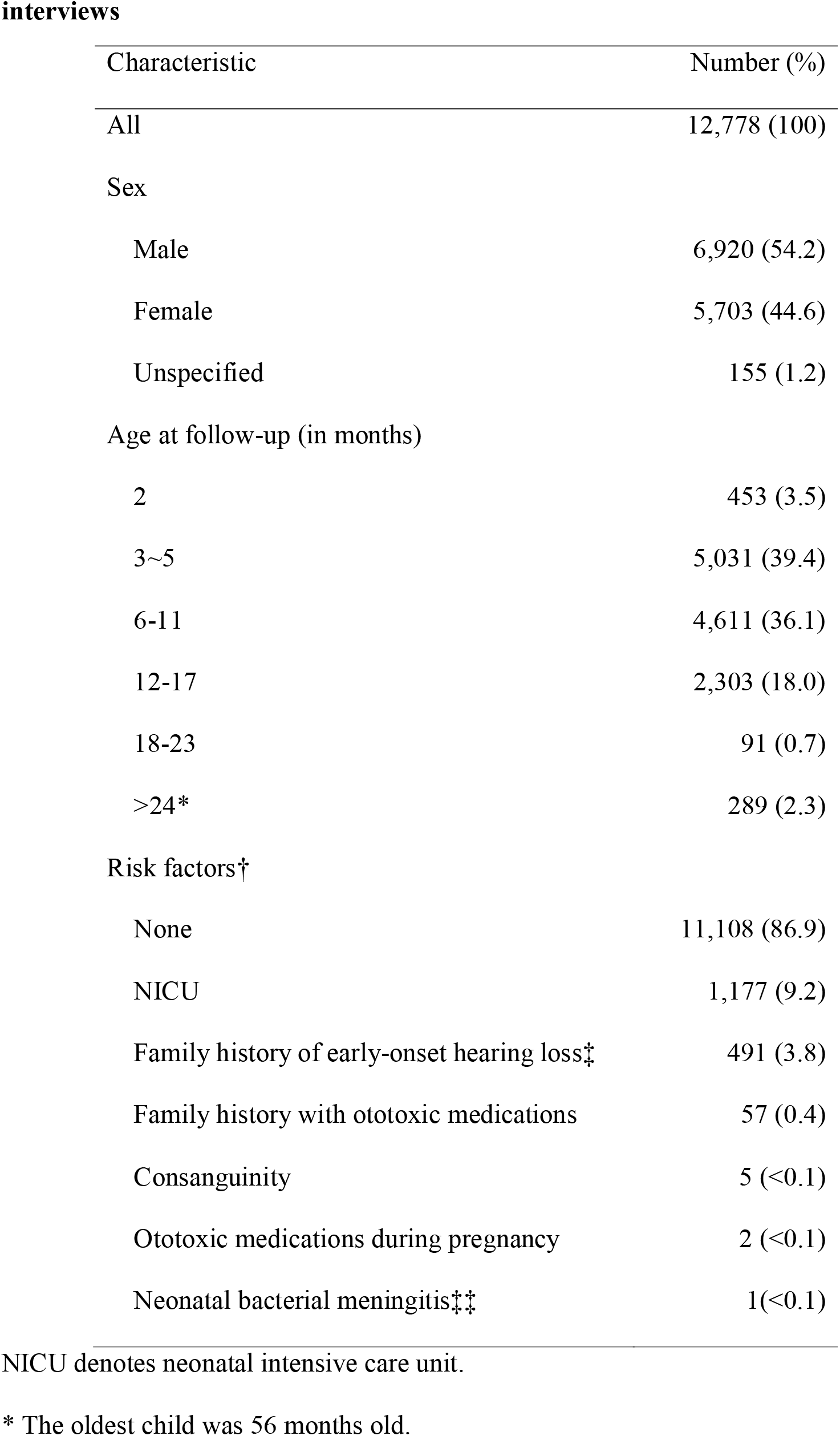
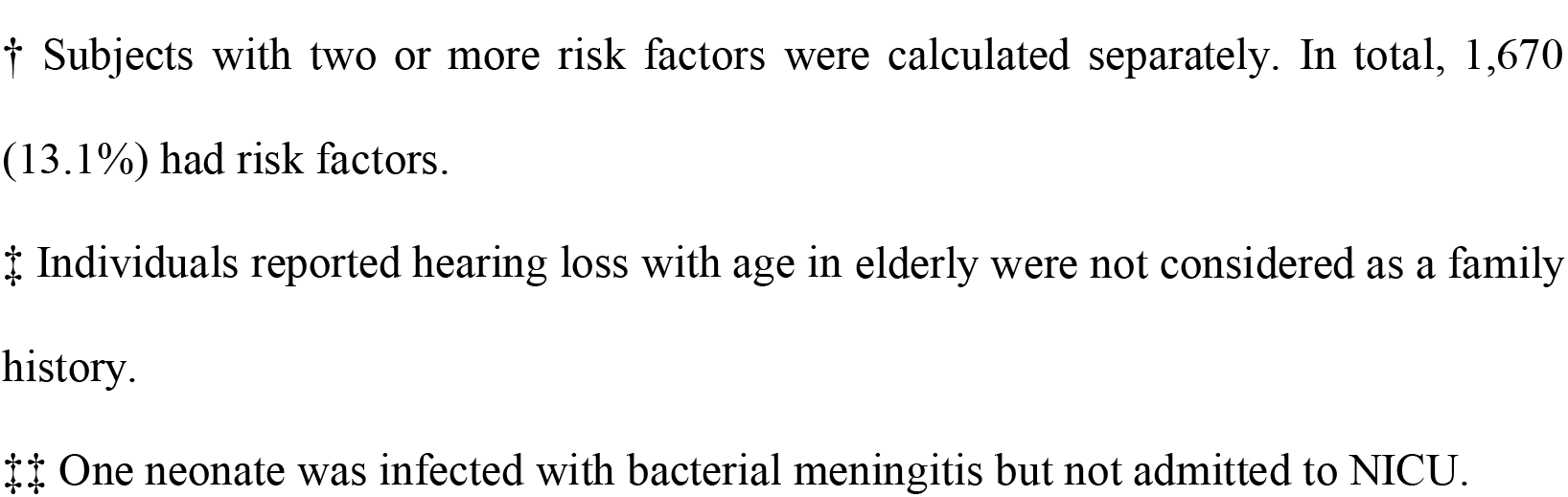
Characteristics of the 12,778 subjects who participated in follow-up.

| Characteristic | Number (%) |
| --- | --- |
| All | 12,778 (100) |
| Sex |  |
| Male | 6,920 (54.2) |
| Female | 5,703 (44.6) |
| Unspecified | 155 (1.2) |
| Age at follow-up (in months) |  |
| 2 | 453 (3.5) |
| 3~5 | 5,031 (39.4) |
| 6-11 | 4,611 (36.1) |
| 12-17 | 2,303 (18.0) |
| 18-23 | 91 (0.7) |
| >24* | 289 (2.3) |
| Risk factors† |  |
| None | 11,108 (86.9) |
| NICU | 1,177 (9.2) |
| Family history of early-onset hearing loss‡ | 491 (3.8) |
| Family history with ototoxic medications | 57 (0.4) |
| Consanguinity | 5 (<0.1) |
| Ototoxic medications during pregnancy | 2 (<0.1) |
| Neonatal bacterial meningitis‡‡ | 1(<0.1) |

NICU denotes neonatal intensive care unit.
\* The oldest child was 56 months old.
† Subjects with two or more risk factors were calculated separately. In total, 1,670 (13.1%) had risk factors.
‡ Individuals reported hearing loss with age in elderly were not considered as a family history.
‡‡ One neonate was infected with bacterial meningitis but not admitted to NICU.

### Benefits of Genetic screening

#### At-risk for ototoxicity

Genetic screening identified 2,638 (0.23%) subjects predisposed to ototoxic hearing loss. Follow-up investigations revealed 57 subjects with a self-reported family history of drug-induced hearing loss (Table 1). Of these, 41 subjects had *MT-RNR1* variants, and 98% (40/41) had a family history consistent with maternal inheritance, which was significantly enriched compared to those without *MT-RNR1* variants (5/16, 31%) (p<0.001) (**Supplementary Table S3** online).

#### Positive Predictive Values

We compared hearing screening results among groups stratified by genetic screening results (Figure 2). The refer rate of the initial hearing screening was 69% (77/112, 95% confidence interval[CI], 59 to 77) in the positive group, which was significantly higher than 5.2% (598/11591, 95% CI, 4.8 to 5.6, P<0.001) and 2.8% (30/1075, 95% CI, 1.9 to 4.0, P<0.001) in inconclusive and at-risk groups, respectively. The refer rate of the 42-day re-test was still higher in the positive group (86%, 49/57, 95% CI, 74 to 94) than in both inconclusive (13%, 61/462, 95% CI, 10 to 17, P<0.001) and at-risk (23%, 6/26, 95% CI, 9 to 44, P<0.001) groups, respectively.

**Figure 2.**
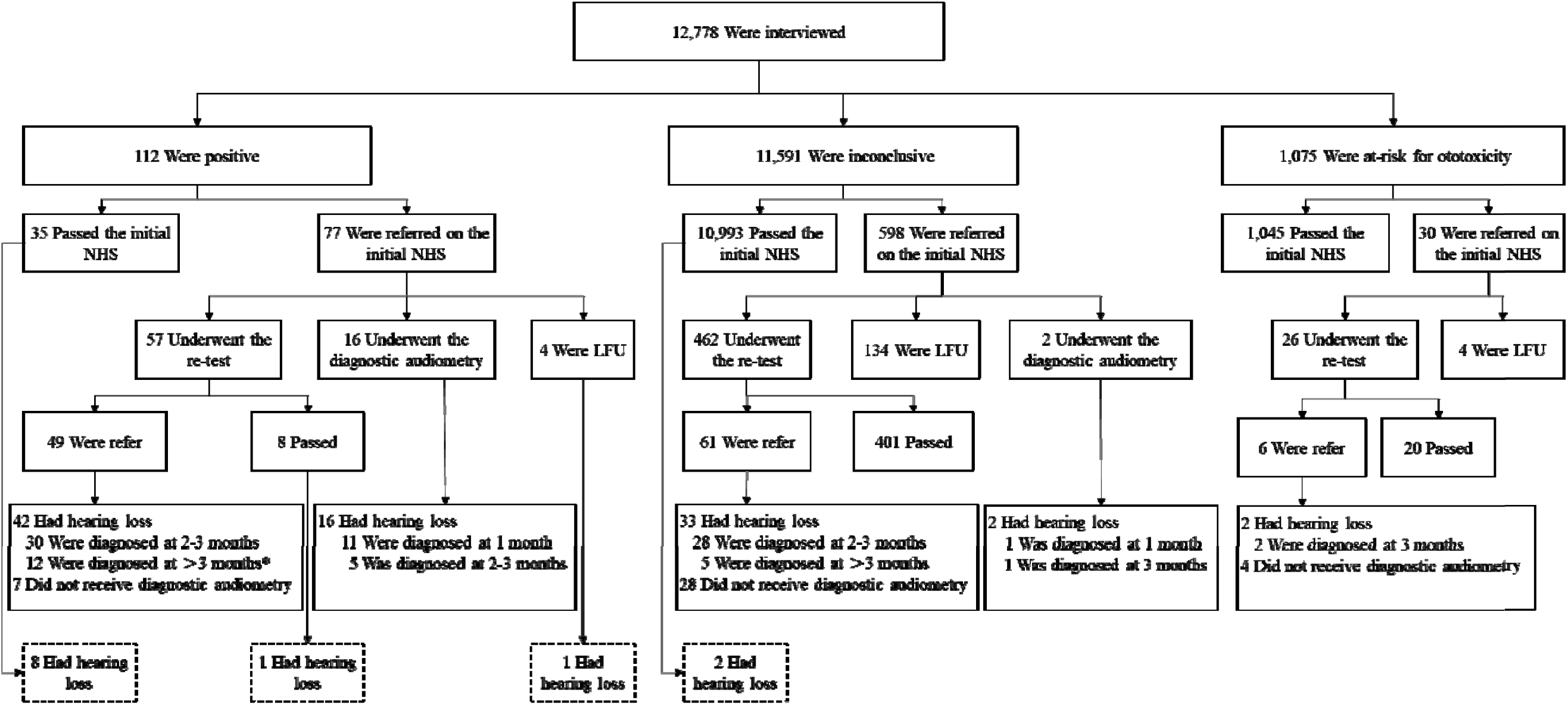
Conventional NHS results of 12,778 newborns with a non-negative genetic screening result who participated in follow-up interviews. NHS denotes newborn hearing screening. LFU denotes loss to follow-up. Dotted box denotes subjects with hearing loss “missed” by the conventional NHS program. * Two subjects (aged 4 and 5 months, respectively) who had not had diagnostic audiometry but who were behaviorally diagn severe/profound hearing loss were categorized in this group.

Overall, of those referred from the initial hearing screening, 46/77 (60%, 95% CI, 48 to 71) in the positve group were diagnosed with hearing loss by three months of age, which was significantly higher than rates in both inconclusive (5.0%, 30/598, 95% CI, 3.4 to 7.1, P<0.001) and at-risk (7%, 2/30, 95% CI, 0.8 to 22, P<0.001) groups (Figure 2). Therefore, a positive genetic screening result indicated the highest PPV of the initial inpatient NHS.

#### Lost to Follow Up

Subjects with different genetic results were compared to evaluate the impact of genetic screening on LFU. The rate of LFU was significantly lower in the positive group (5%, 4/77, 95% CI, 1 to 13) than that in the inconclusive group (22%, 134/598, 95% CI, 19 to 26, P<0.001), but there was no significant difference between the at-risk and inconclusive groups (13% (4/30) vs. 22%, P = 0.24).

#### Hearing loss cases “Missed” by hearing screening

Among the cohort eligible for the primary analysis, 107 subjects were reported to have hearing loss (Figure 1) by the time of follow-up interviews (mean age = 9.5±8.7, median age = 7 months, range 2–56 months). Of these, only 95 subjects were diagnosed following the protocol of the conventional NHS program (Figure 2). The remaining 12 subjects (ten positive and two inconclusive) fell into three categories: ten passed the initial hearing screening, one passed the 42-day re-test, and one was LFU (Table 2). None of the 12 subjects could be identified by risk-factor-indicated audiologic monitoring.

**Table 2.**
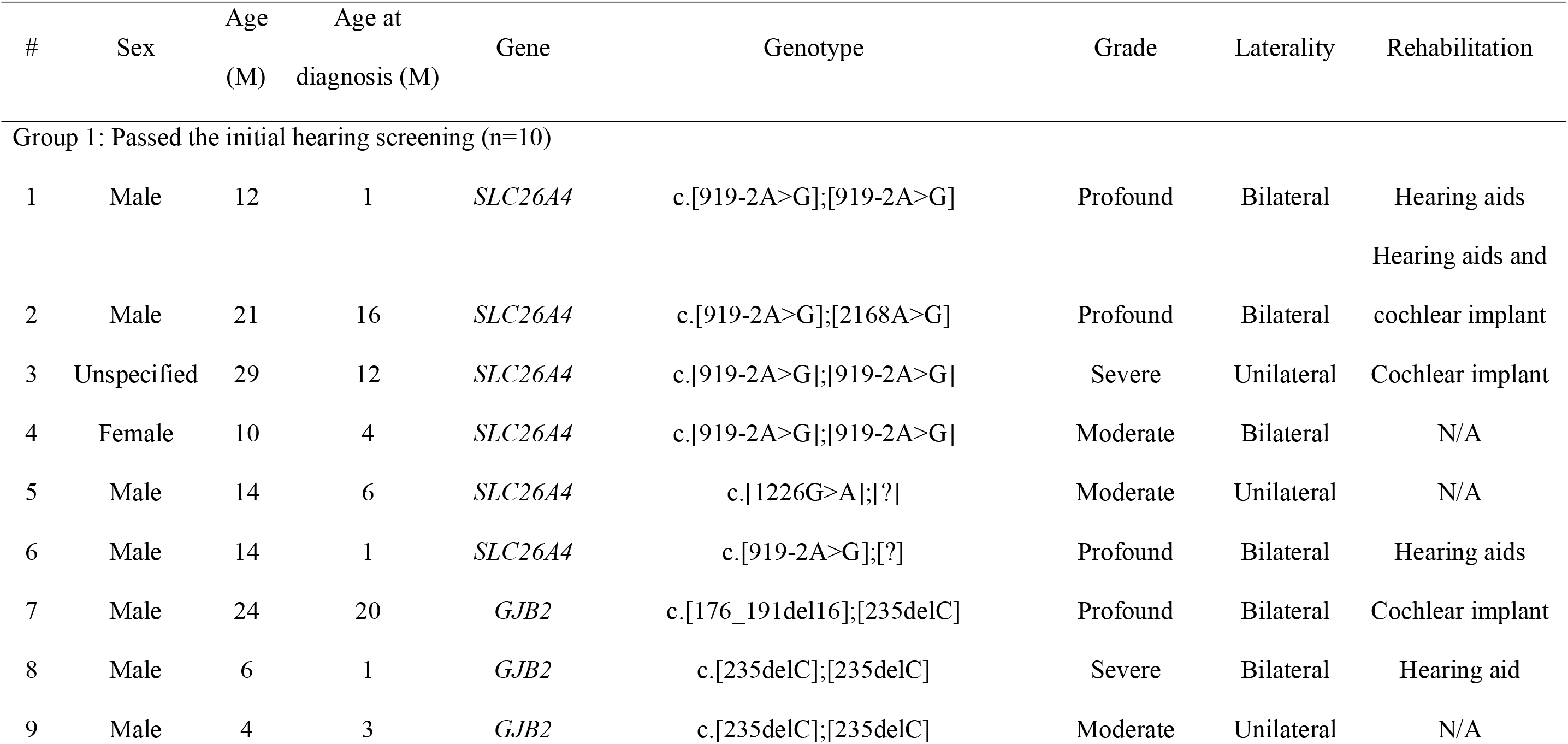
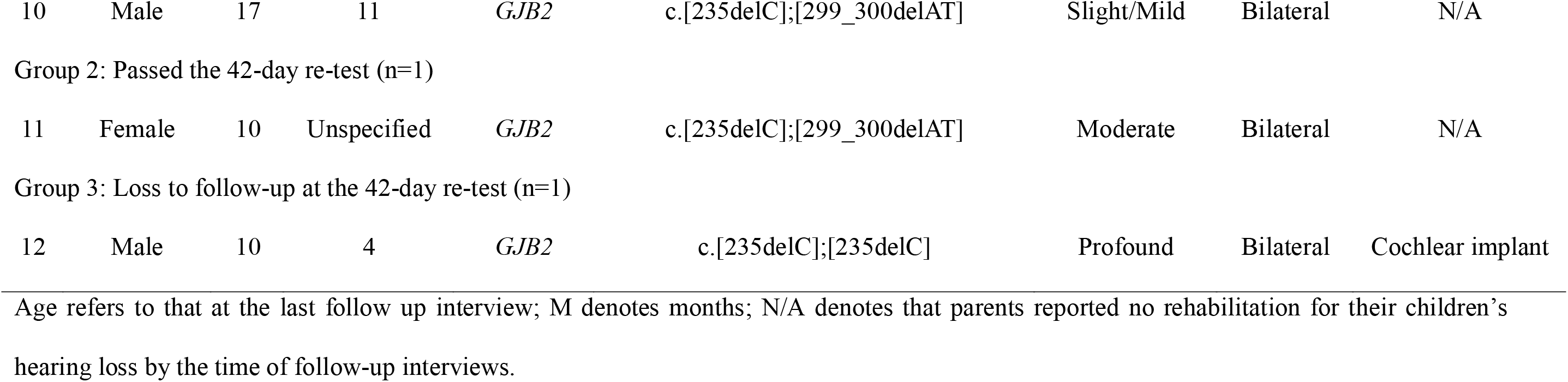
Characteristics of subjects “missed” by the conventional newborn hearing screening program who developed hearing loss.

Because hearing loss is highly penetrant in individuals with biallelic pathogenic variants in *GJB2* or *SLC26A4* on our screening panel, all 112 subjects with a positive genetic result are expected to develop hearing loss eventually. Of these subjects, 47 (42%, 95% CI, 33 to 52) were “missed” by the conventional NHS program (35 passed the initial inpatient NHS, 8 passed the 42-day re-test, and 4 were LFU), suggesting a 72% (47/65) potential improvement of detection yield in the positive group. Notably, only 10 of 47 subjects would be detected by risk-factor-indicated audiologic monitoring.

### Genotype and phenotype correlation

The secondary analysis was focused on the genotype and phenotype correlation in 107 subjects confirmed to have hearing loss by the time of follow-up interviews (Table 3).

**Table 3.**
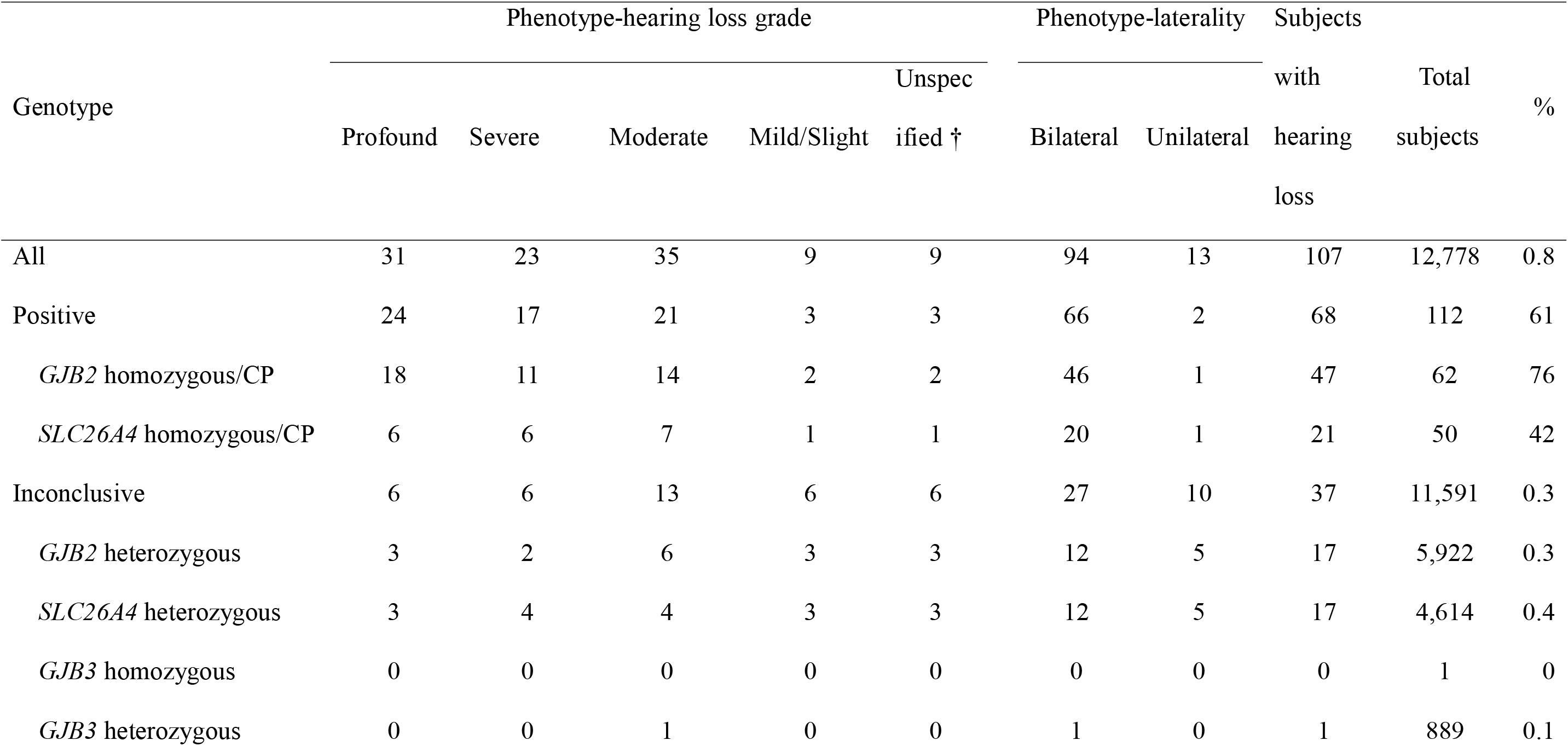
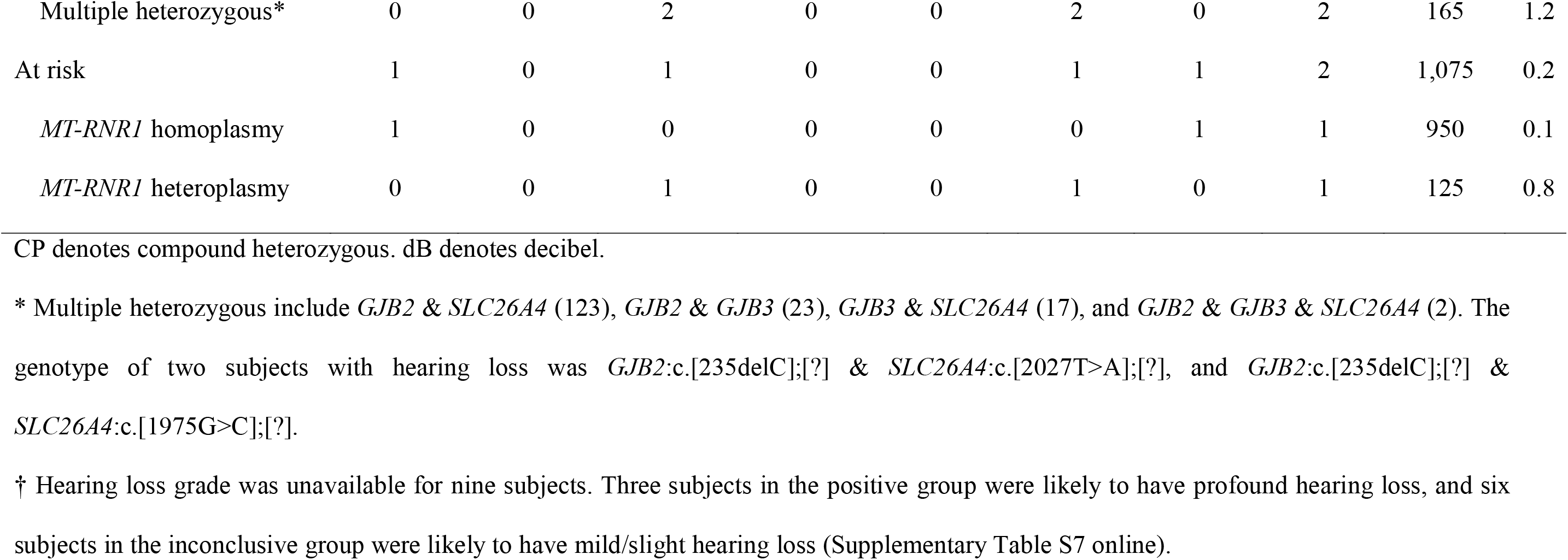
Genotype-phenotype correlation for the 107 subjects with hearing loss.

Of 112 subjects in the positive group, 68 (61%, 95% CI, 51 to 70) were reported to have hearing loss. Mean ages for *GJB2* and *SLC26A4* positive subgroups were not significantly different (10.4±9.6 vs. 11.0±9.3 months, P = 0.60). However, 76% (95% CI, 63 to 86) of subjects with a positive *GJB2* result developed hearing loss, whereas the rate was 42% (95% CI, 28 to 57) in the *SLC26A4* positive subgroup (P<0.001). Both subgroups had predominantly bilateral hearing loss (46/47, 98% and 20/21, 95%, respectively). Moreover, subjects in the positive group were more likely to have profound or severe hearing loss (60%) compared to those in the inconclusive group (32%) (P = 0.006) (Table 3).

Of 11,591 subjects in the inconclusive group, 37 (0.3%, 95% CI, 0.2 to 0.4) were reported to have hearing loss, slightly higher than but not significantly different from the population prevalence. Of one *GJB3* homozygote (age 13 months) and 889 *GJB3* heterozygotes (mean age = 8.1±5.5 months, median age = 7 months, range 2 to 44 months), only one *GJB3* heterozygote was diagnosed with hearing loss (**Supplementary Table S4 online**). None of the 25 *GJB2/GJB3* double heterozygotes (mean age = 7.6±3.9 months) reported hearing loss at the time of the last follow-up interviews (**Supplementary Table S5** online). Hearing loss family history was not enriched in subjects with *GJB3* variants (3.1%, 28/915) over those without (4.1%, 463/11372, P = 0.23).

Of 1,075 subjects in the at-risk group, only two had hearing loss (**Supplementary Table S6** online). None of them was reported to have had any exposure to ototoxic drugs; therefore, the mitochondrial variant identified was unlikely the cause of hearing loss in either case.

## Discussion

Here we report results of a large-scale genetic screening program in China. The goal of genetic screening was not to diagnose all hereditary deafness but rather to supplement conventional newborn hearing screening to ensure that children with hearing loss receive timely identification and intervention. We chose a limited screening panel for population genetic screening to fulfill the purpose, because it has the advantages of low cost, fast turnaround time, and easy interpretation and low uncertainty of results over comprehensive diagnostic testing methods. Even with just the limited panel, our study has already demonstrated significant benefits of genetic screening.

Our primary analysis demonstrates that genetic screening informed 0.23% of newborns and their maternal family members of susceptibility to potentially preventable ototoxicity due to *MT-RNR1* variants by avoiding exposure to aminoglycoside antibiotics. Notably, all subjects from 41 families with a self-reported family history of drug-induced hearing loss had avoided aminoglycoside antibiotics and none of them developed hearing loss thanks to the knowledge and genetic counseling.

Incorporating genetic screening into the NHS program can identify additional newborns with hearing loss at an earlier age.^13–15^ Our genetic screening platform detected 72% more children at two to four weeks of age with reported or presymptomatic hearing loss in the positive group, who were “missed” by hearing screening due to delayed onset or LFU. Given the 0.03% genetically positive rate, the 58% diagnostic yield of hearing screening in the positive group, and the 0.1–0.3% prevalence of congenital hearing loss in the general population, our genetic screening is estimated to detect 6–17% of all newborns with hearing loss diagnosed by hearing screening and another 4–13% that would have been “missed”. Hence, the benefit of genetic screening is remarkable, compared to a <1% additional yield by risk-factor-indicated audiologic monitoring recommended by the Joint Committee on Infant Hearing.^16–18^

Low PPV of the initial hearing screening step is a limitation of the conventional NHS program, because it brings unnecessary parental anxiety^19^ and increases healthcare cost for follow-up evaluations.^20^ Studies have reported PPVs of 2.2% to 15% in clinical practice.^5,6^ We observed similar low PPVs in the inconclusive (5%) and at-risk (7%) groups, which should be representative of that in the general population. However, the PPV was significantly higher in the positive group (60% by three months). The theoretical PPV is the penetrance of hearing loss in the positive group, which would approach 100%.

LFU undermines the clinical effectiveness of NHS programs.^21^ We report that the LFU rate was significantly lower in the positive group (5%) than in the inconclusive group (22%) who were expected to be representative of the general population. Notably, one of four subjects in the positive group who were LFU by the conventional NHS program developed hearing loss by the time of the last interview, and the remaining were predicted to be likely to develop hearing loss later in life and aware of the situation thanks to genetic screening. Hence, incorporating genetic screening into the NHS program would not only significantly decrease LFU but also prompt active surveillance of subjects with a positive genetic result. The positive group is indeed the target population who can benefit the most from early intervention for hearing loss.

Our secondary analysis focused on genotype-phenotype correlations. By the time of follow-up interviews, 44 subjects in the positive group aged 3 to 38 months were reported to be hearing based on their latest audiologic evaluation or behavioral observation. They are expected to develop hearing loss eventually because the penetrance of hearing loss in individuals with biallelic pathogenic variants in *GJB2* or *SLC26A4* on our screening panel is nearly complete taking into account variable age of onset.^22–24^ Long-term longitudinal follow-up is necessary to document their hearing status. In the inconclusive group, 37 subjects were reported to have hearing loss. Of note, *GJB2* or *SLC26A4* heterozygotes might have a second pathogenic allele in the same gene not screened by our panel. Identifying the second allele in these subjects would not only provide an etiologic diagnosis and be of value in genetic counseling, but also inform future panel design of recurrent alleles with relatively high frequencies. Nevertheless, our data did not provide evidence to support the pathogenicity of *GJB3* variants in autosomal recessive, dominant, or digenic hearing loss, consistent with a recent study.^25^ Although their roles in late-onset recessive or digenic hearing loss cannot be determined, the lack of enrichment of hearing loss family history in subjects with these variants indicates they do not cause late-onset dominant hearing loss.

The benefits of genetic screening could be enhanced further by optimizing ethnicity-specific panels and increasing the number of variants without increasing the cost and turnaround time. The current screening panel was designed in 2011 when large population databases with ethnicity-specific allele frequencies were unavailable. We now observe *GJB2*:c.35delG, *GJB2*:c.167delT, and *SLC26A4*:c.2162C>T with high frequencies in European, Jewish, and Latino populations, respectively, in the Genome Aggregation Database,^26^ but very low or even undetected in our Chinese study population (**Supplementary Table S1** online). Replacing these three variants and the *GJB3* variants with other relatively common pathogenic variants in the Chinese population is anticipated to improve the panel.

The clinical practice upon receiving genetic screening results varied in different hospitals, which could be further optimized. Sixteen subjects in the positive group referred from the initial hearing screening underwent the diagnostic audiologic evaluation directly. Eleven of them were diagnosed with hearing loss at one month of age, earlier than the average time to diagnosis. In addition, 35 subjects in the positive group developed hearing loss by three months of age. Our data suggest that subjects should schedule an audiologic evaluation directly upon receipt of a positive genetic result. As a comparison, although very few subjects in the inconclusive group underwent the diagnostic audiologic evaluation directly and received a earlier diagnosis as well, only 6% (33/598) of subjects who were referred from the initial hearing screening developed hearing loss by the time of the last interviews. Therefore, to avoid unnecessary anxieties and medical costs, we do not recommend that subjects with an inconclusive genetic screening result bypass the re-test. Nevertheless, our findings call for professional guidelines for the global implementation of concurrent NHS programs.

Strengths of this study are a large sample size and outcome assessment with sufficient power to demonstrate the benefits of genetic screening. It provides evidence and experience for other countries, hospitals, and laboratories to implement such strategies to meet the urgent need of improving NHS. The weaknesses include vague recalls from some interviewees, inability to follow up newborns with negative genetic screening results who did not pass the initial inpatient NHS, and insufficent long-term follow-up to assess intervention outcomes, which warrant further investigations.

In conclusion, this study provides convincing evidence that incorporating a limited genetic screening panel into NHS is an effective strategy to improve NHS programs by providing etiologic diagnoses, identifying hearing loss “missed” by hearing screening, increasing PPV and reducing LFU in the positive group, and alerting at-risk newborns and their maternal relatives of their susceptibility to ototoxicity.

## Acknowledgments

This study was supported by Shenzhen Engineering Laboratory for Innovative Molecular Diagnostics (DRC-SZ[2016]884); by grants (to Dr. Wang) from the National Natural Science Foundation of China (Major Project No.81530032) and the National Key Basic Research Program of China (2014CB943001); by grants (to Drs. Morton and Shen) from the National Institutes of Health/the National Institute on Deafness and Other Communication Disorders (R01DC015052) and (to Dr. Song) from the National Natural Science Foundation of China (Grant No.81600690).

